# Inhibition of KDEL receptors remodels the tumor microenvironment for robust T cell independent tumor regression

**DOI:** 10.1101/2025.08.07.669177

**Authors:** Shakti P Pattanayak, Hong Wang, Belinda Willard, Timothy A. Chan, William C Merrick, Zheng-Rong Lu, Boaz Tirosh

**Affiliations:** Department of Biochemistry, Case Western Reserve University School of Medicine, Cleveland, OH, USA; Department of Biomedical Engineering, Case Western Reserve University School of Medicine, Cleveland, OH, USA; Proteomics and Metabolomics Shared Laboratory Resource, Lerner Research Institute, Cleveland Clinic, Cleveland, OH, USA; Center for Immunotherapy and Precision Immuno-Oncology, Cleveland Clinic, Cleveland, OH, USA

**Author notes:** Send correspondence to: Boaz Tirosh, Case Western Reserve University, Department of Biochemistry, Cleveland, OH, 44106, USA Tl.

**Keywords:** calreticulin, ER stress, immunogenic cell death, innate immunity, immunotherapy, cancer immunity cycle

## Abstract

Tumor immunotherapy is supported by low-grade inflammatory conditions at the microenvironment, triggered by immunogenic cell death (ICD). However, ICD is dampened when tumors acquire resistance, affecting immune recognition. KDEL receptors (KDELRs), through a retrograde Golgi-to-ER transport, prevent spontaneous secretion of KDEL proteins. We report that inhibition of a single KDELR in a minor fraction of tumor cells, primarily KDELR2, provokes robust infiltration of macrophages and neutrophils into the tumor microenvironment, resulting in a complete regression of both immunogenic and non-immunogenic tumors independently of T cells. Importantly, in the course of regression, anti-tumor T cells are primed, conferring protection against a second challenge. Recapitulated by intratumoral delivery of siDKELR2 utilizing lipid nanoparticles, we implicate KDELR2 as a target to unleash an unusual robust innate immune response, which represents a tractable approach to initiate an adaptive response downstream, bypassing conventional ICD-inducing therapies. We propose KDELR targeting as a strategy to improve immunotherapy across tumor types, including “cold” tumors resistant to T cell-based immunotherapies.

## Introduction

The cancer immunity cycle (CIC) model has been evoked to describe the iterative steps by which the anti-cancer immune response is perpetuated and adapts to evolving tumors. A productive CIC is required to achieve a durable immune response and is dependent on the maintenance of inflammatory conditions in the tumor microenvironment (TME), which switches the immune response from tolerogenic to immunogenic (Mellman *et al*, 2023). A major driver of the inflammatory TME is the immunogenic cell death (ICD) of tumor cells, which is a form of cell death that attracts macrophages and dendritic cells (DCs) into the tumor tissue (Becker *et al*, 2002). Inflammatory conditions *in situ* facilitate the productive activation of effector immune cells, primarily T cells and natural killer (NK) cells (Kepp *et al*, 2019), and are therefore highly desired for cancer immunotherapy (Chen *et al*, 2023; Jiang *et al*, 2022). ICD is generated when cell death is accompanied by cellular stress, typically induced by harsh cancer therapies, such as chemotherapy (Vacchelli *et al*, 2014), radiotherapy (Wang *et al*, 2023a), and photodynamic therapy (Hao *et al*, 2022). However, when tumors gain resistance to therapy, stress conditions are mitigated by adaptation, causing ICD to subside and inflammatory conditions in the TME to decay. This allows tumors to overcome immune recognition and regain their growth.

Inflammatory conditions that develop during ICD are due to *in situ* release of damage-associated molecular patterns (DAMPs), which are immune stimulatory molecules that reside intracellularly and are secreted during cell death(Tang *et al*, 2024). The number of confirmed DAMPs has increased in recent years, ranging from small molecules to proteins, glycoproteins, and proteoglycans (Hayashi *et al*, 2020; Kepp *et al*, 2014). A prominent DAMP is calreticulin (Kepp *et al*, 2020), a highly abundant endoplasmic reticulum (ER) luminal chaperone protein encoded by the ER retention C-terminal sequence KDEL. During ICD, a small fraction of calreticulin is secreted and/or enriched at the cell surface, termed ectoCalreticulin, where it is recognized by natural killer (NK) cells (Sen Santara *et al*, 2023) and macrophages (Ogden *et al*, 2001). Of note, calreticulin is one of many KDEL proteins, all of which rely on this sequence for ER retention.

The underlying mechanism of KDEL-mediated ER retention is evolutionarily conserved from yeast to mammals, involving retrograde transport from the Golgi back to the ER of leaked KDEL proteins by three seven transmembrane KDEL receptors (KDELR1-3). KDELRs utilize a sophisticated catch-and-release mechanism that exploits the differences in pH between the Golgi and ER to bind the KDEL protein in the Golgi, traffic to the ER, and release the cargo, allowing empty KDELRs to return to the Golgi to resume their gatekeeper function (Newstead & Barr, 2020). Although different functions have been attributed to each of the KDELR isoforms, most likely related to their different abilities to signal as GPCR receptors, all three isoforms participate in preventing the secretion of KDEL proteins (Newstead & Barr, 2020). This quality control mechanism is sensitive to stress conditions. Treatment with chemicals that perturb protein folding in the ER, such as thapsigargin (Tg), or hypoxia, results in KDEL protein secretion (Han *et al*, 2019). Moreover, KDELRs translocate to the cell surface under stress conditions, further promoting the secretion of KDEL proteins (Becker *et al*, 2016). Studies on different cell types have shown that the quality control mechanism is near or at saturation. Knockdown of a single KDELR, primarily KDELR2, is sufficient to induce the secretion of KDEL-decorated proteins in a manner that synergizes with ER stress conditions (Trychta *et al*, 2018).

Since the mechanism of intracellular retention of KDEL proteins is common to all KDEL proteins, it is expected that multiple KDEL proteins, not only calreticulin, are displaced *en bloc* from the ER during ICD. Additional KDEL proteins, such as protein disulfide isomerases (PDI), Bip, and gp96, have been reported to translocate to the surface under cellular stress conditions (Cifric *et al*, 2024; Kaiser *et al*, 2007; Soares Moretti & Martins Laurindo, 2017; Vabulas *et al*, 2002). Bioinformatic studies have shown a correlation between high KDELR expression and poor prognosis in various tumor types (Cela *et al*, 2022) and a reduced presence of immune cells in the TME (Li *et al*, 2024). Direct investigation of KDELRs as modulators of the TME and immune recognition in cancer has not been documented. The current study provides evidence that the inhibition of KDELR2 in a small fraction of tumor cells is sufficient to attract massive amounts of innate immune cells, primarily macrophages, into the TME. Under these conditions, tumor regression and systemic protection are generated in the absence of any other intervention. We propose that targeted KDELR suppression in tumor cells could be a strategy to initiate ICI therapy and enhance tumor immunotherapy.

## Results

### Suppression of KDELRs is sufficient to confer secretion of KDEL proteins

To study the effect of KDELRs on TME, we used the antigenic colon cancer cell line MC38, which was isolated from female C57BL/6 mice (Corbett *et al*, 1975). Subcutaneously implanted MC38 cells grow rapidly in naïve C57BL/6 mice. However, growth is inhibited by vaccination (McLaughlin *et al*, 1997) and further inhibited by the administration of immune checkpoint inhibitors (Allard *et al*, 2013). Because MC38 is responsive to photodynamic therapy (Nagaya *et al*, 2019), we reasoned that MC38 growth in C57BL/6 hosts would be a suitable model to study the role of KDELR-KDEL quality control in cancer immunotherapy. To assess which of the three KDELRs is the most dominant in the retention of KDEL proteins, we stably expressed nanoluc luciferase in MC38 cells, equipped with a signal peptide at the N-terminus and a KDEL sequence at the C-terminus. We then introduced a Tet-On shRNA sequence for each KDELRs by lentiviral infection (**Fig. 1A**). Since KDELRs are highly homologous to each other in the amino acid sequence, commercial KDELR antibodies do not distinguish between the different isoforms. Therefore, we used qPCR analyses to confirm shRNA activity 24 h after the addition of doxycycline (DOX) to the medium. As expected, each shRNA was specific for the respective KDELR isoform (**Fig. 1B**). Luciferase activity was measured in the supernatants. DOX-induced silencing of KDELR2 resulted in the highest increase in luciferase activity in the medium. Silencing of KDELR1 reduced luciferase activity in the medium, while the silencing of KDELR3 did not result in any significant effect (**Fig. 1C**). These results were congruent with those observed in SH-SY5Y cells transfected with siRNA (Trychta *et al*., 2018). Calreticulin and other ER proteins adhere to the cell surface upon secretion. Flow cytometry analysis of Tet-On shKDELR2 cells for ectoCalreticulin showed an increase following the addition of DOX (**Fig. 1D**, quantified in 1E). Because stress conditions promote the accumulation of ectoCalreticulin, ectoBip, and surface PDIs (Gutierrez & Simmen, 2014), we examined whether KDELR2 suppression can further promote the secretion of KDEL proteins under stress conditions. Cooperation between hypoxic conditions and KDELR2 suppression in promoting surface calreticulin was observed (**Fig. 1D**, quantified in 1E), accompanied by higher secretion of nanoluc-KDEL. A similar effect was observed for ER stress induced by Tg (Fig. S1). These data suggest that *in vivo*, during tumorigenesis, when multiple stress conditions develop simultaneously, silencing of KDELR2 should further intensify the release of DAMPs.

**Figure 1:**
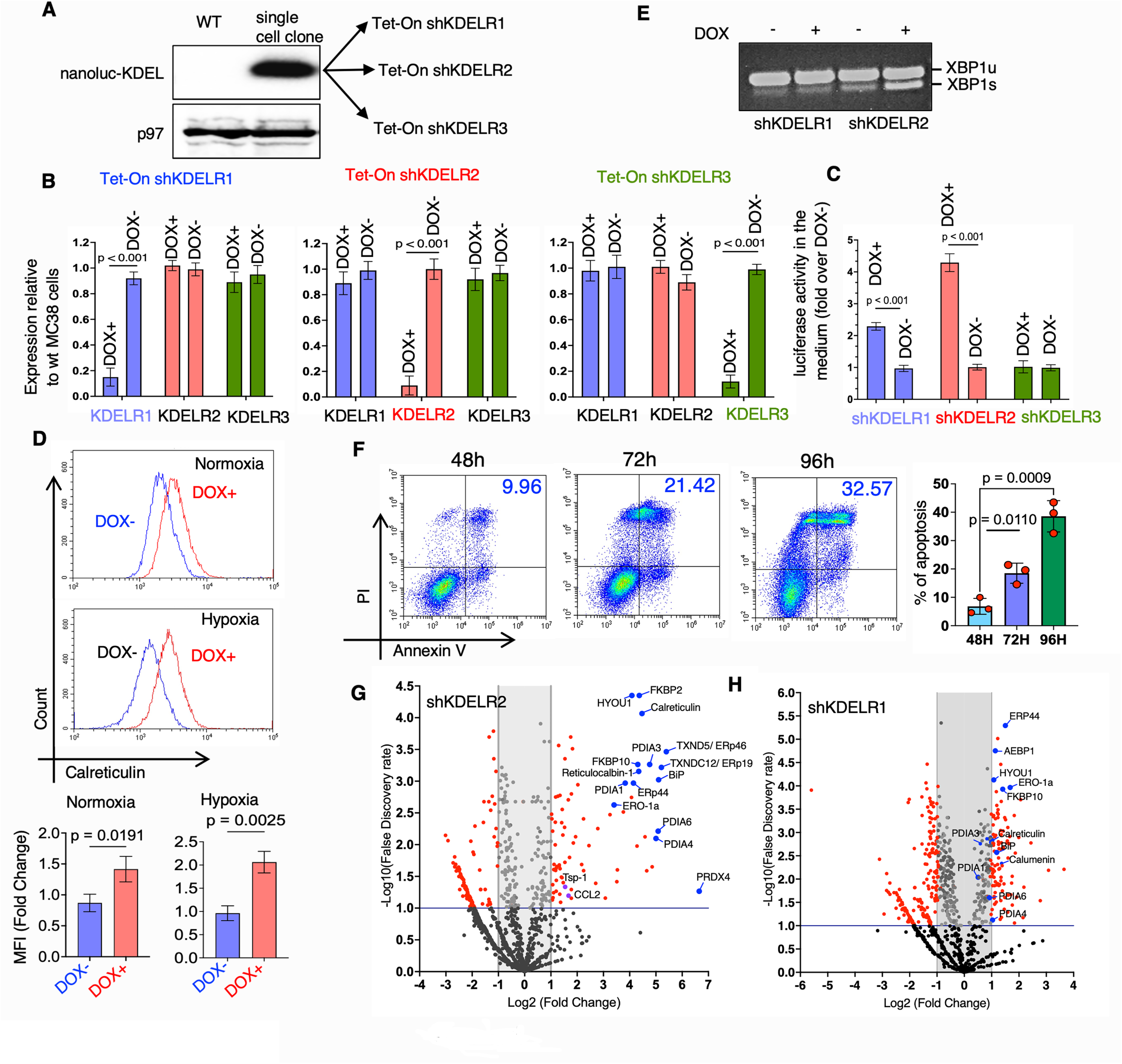
Silencing of KDELR2 is sufficient to induce secretion of secretory KDEL and non-KDEL proteins. **A**. MC38 cells were stably transfected with nanoluc-KDEL. Cells were then infected with lentiviruses that encode Tet-On shRNA for KDELR1, KDELR2 or KDELR3. **B**. qPCR analysis 24 h after the addition of DOX (10 µM) confirmed activity and specificity of the shRNA sequences. Shown is the average and SD of three technical repetitions. **C**. Analysis of supernatant luciferase activity following 24 h incubation with DOX. Activity was normalized to activity without DOX. Shown is the average and SD of three technical repetitions. **D**. Flow cytometry for ectoCalreticulin following 24 h of incubation with DOX under normoxic and hypoxic conditions. Quantified in **E**. **F**. RT-PCR analysis of XBP1 mRNA splicing as evidence of ER stress in after 24 h of DOX treatment of shKDELR1 and shKDELR2 cells. **G.** Flow cytometry analysis for apoptosis of MC38 Tet-On shKDELR2 cells at times after the addition of DOX. **H** and **I**. Volcano plots for enrichment of proteins in the supernatats of MC38 Tet-On shKDELR2 or MC38 Tet-On shKDELR1 after a treatment for 24 h with DOX.

We reasoned that by compromising KDELR2 expression, protein folding in the ER would be compromised, instigating the adaptive unfolded protein response (UPR). A hallmark of the UPR is the splicing of XBP-1 mRNA by the UPR transducer IRE1 (Yoshida *et al*, 2001). RT-PCR for XBP1 mRNA demonstrated enhanced splicing after DOX addition for 24 h in Tet-On shKDELR2 cells (**Fig. 1F**). Since unmitigated conditions of ER stress confer apoptosis (Szegezdi *et al*, 2006), we analyzed the level of apoptosis by flow cytometry using Annexin V and propidium iodide (PI) over time. Growth retardation and increased apoptosis were detected after 48 h of DOX treatment and increased with treatment duration (Fig. S1, **Fig. 1G**), suggesting that the constant suppression of KDELR2 confers irreparable ER stress. However, this effect developed over a few days. To analyze the effect of KDELR suppression on the cellular secretome, we collected the supernatants for mass spectrometry-based proteomics. To avoid confounding factors related to serum proteins, cells were cultured for 24 h in OptiMem medium without serum, supplemented with insulin/transferrin/selenium with and without DOX. At this time point, the viability was minimally affected. Since the suppression of KDELR3 did not affect the secretion of nanoluc-KDEL, we only analyzed MC38 cells transduced with shRNA for either KDELR1 or KDELR2. In accordance with the luciferase activity measurements, silencing of KDELR2 resulted in a 20-30 fold increase in the level of KDEL proteins, labeled in blue, such as Bip, PDIA6, PDIA4, and PDIA3 (**Fig. 1H**). KDELR1 suppression was less pronounced (**Fig. 1I**). Notably, the secretome does not include surface-attached proteins, thus underestimating the total leakage of secretory proteins. Notably, non-KDEL proteins are enriched in the supernatants, some with immunomodulatory functions, such as thrombospondin-1 (TSP-1) (Lopez-Dee *et al*, 2011) and chemokine C-C motif ligand 2 (CCL2) (Lin *et al*, 2022).

### Suppression of KDELR2 expression in a small fraction of tumor cells results in infiltration of innate immune cells and T cell-independent tumor regression

To address the role of KDELR2 in tumorigenicity, we subcutaneously challenged C57BL/6 mice in the left flank with wild-type MC38 cells and in the right flank with Tet-On shKDELR2 MC38 cells. Twenty-four hours before the challenge, the mice were fed either a normal chow diet (DOX-) or a doxycycline-supplemented diet (DOX+). Tumor size was assessed throughout the duration of the experiment using calipers. While WT MC38 cells established tumors in both DOX-and DOX+ fed mice, Tet-On shKDELR2 cells established tumors when normal chow was used at similar growth kinetics to wt. MC38 Tet-On shKDELR2 did not establish tumors when the mice were fed with DOX prior to challenge (Fig. S2). We then tested the effect of KDELR2 suppression once tumors are established. Mice were challenged in both flanks with wt MC38 and Tet-On shKDELR2 cells. When either of the tumors reached approximately 300 mm^3^, the diet was switched to DOX+. While wt MC38 tumors continued to grow, Tet-On shKDELR2 tumors regressed and completely disappeared within two weeks (**Fig. 2A**), consistent with the impeded growth of Tet-On cells *in vitro* when exposed to DOX. We then tested whether the suppression of KDELR2 in a small fraction of tumor cells could affect the tumorigenicity of the majority of WT MC38 cells. To this end, we mixed MC38 with Tet-On shKDELR2 cells prior to inoculation at a 10:1 ratio (denoted as “MIX 10:1”). C57BL/6 mice were challenged with MC38 (left flank) and MIX 10:1 (right flank). When tumors reached approximately 200 mm^3^, the mice were fed a DOX diet. We observed that MIX 10:1 tumors continued to grow for a few days and then regressed. After 28 days, MIX 10:1 was hardly detected, while MC38 tumors continued to grow (**Fig. 2B**). We assumed that the regression was dependent on T cells, since MC38 is antigenic and susceptible to T cell-mediated responses (Denis *et al*, 2022). Therefore, we repeated the experiment in nude mice lacking T-cells. Similar to C57BL/6 mice, under the DOX-supplemented diet, MIX 10:1 tumors underwent regression (**Fig. 2C**), indicating that the regression does not require T cells.

**Figure 2:**
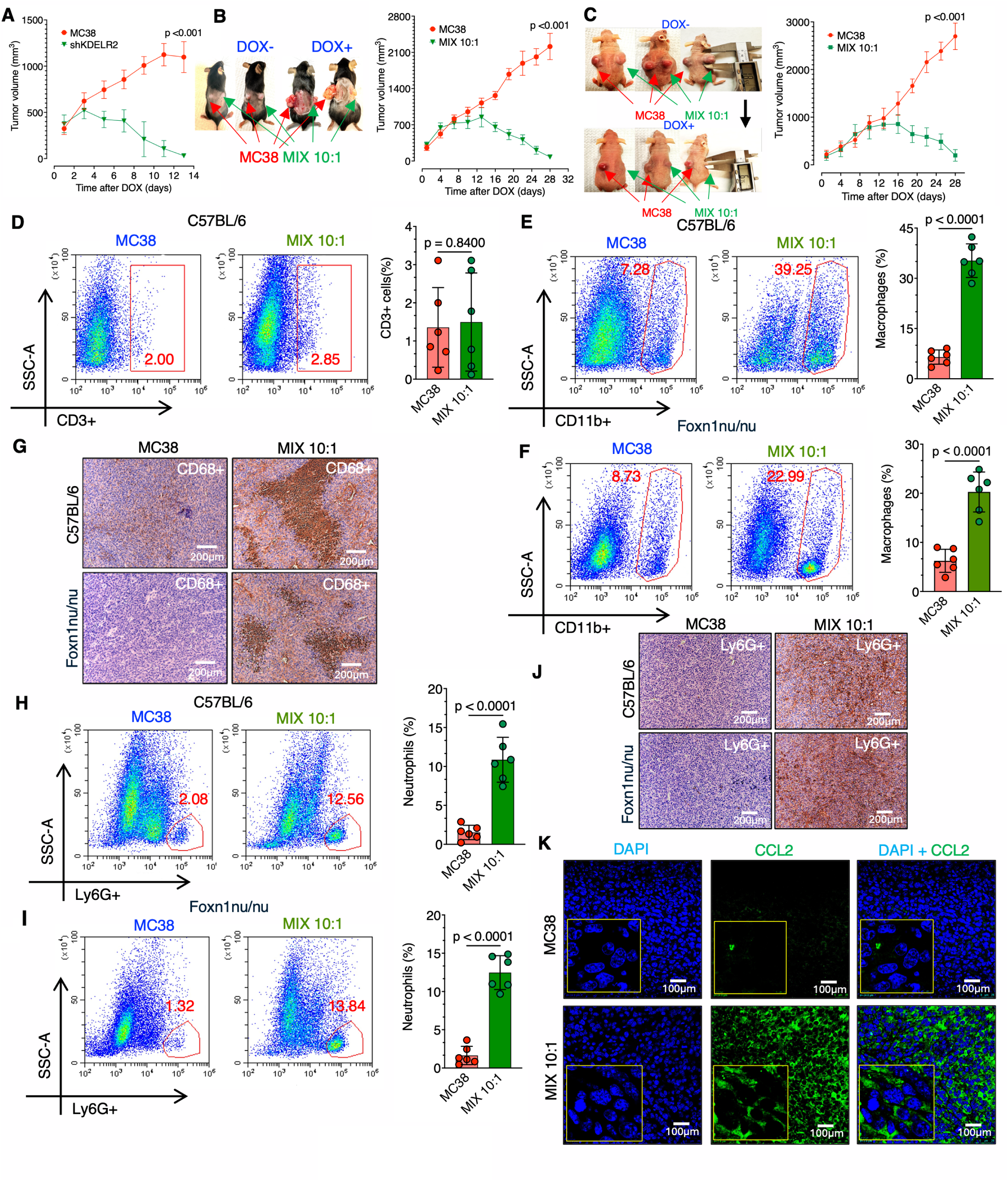
Silencing of KDELR2 in a small percentage of tumor cells confers regression in a T cell independent manner associated with the infitration of macrophages and neutrophils into the TME. **A.** Cohorts of 6 C57BL/6 animals were implanted s.c. with MC38 cells on the left and MC38 with Tet-On shKDELR2 on the right. **A.** Mice were put on regular chow and moved to DOX supplemented diet when tumors exceeded 300 mm^3^. Tumor size was quantified by a caliper and quantified. **B.** Tumor growth as in A, but on the right flank a MIX 10:1 was implanted. **C**. Tumor growth of MC38 on the right flank and MIX 10:1 tumors on the left in nude mice. **D**. Shown are representative flow cytometry analyses for T cells and quantification from 5 different tumors. **E** and **F**. Representative flow cytometry analyses for macrophages in WT and MIX 10:1 tumors grown in C57BL/6 (**E**) and nude mice (**F**). Qunatification of five tumors is shown on the right. **G**. Immunohistochemistry analysis for macrophages demonstrating clusters in the MIX 10:1 tumors 10 days after placing the mice on a DOX diet. **H** and **I**. Representative flow cytometry analyses for neutrophils in WT and MIX 10:1 tumors grown in C57BL/6 (**H**) and nude mice (**I**). Quantification is shown on the right. **J**. Immunohistochemistry analysis for neutrophils in the MIX 10:1 tumors 10 days after placing the mice on a DOX diet. **K**. Immunohistochemistry analysis for CCL2 in wt and MIX 10:1 tumors 10 days after placing the mice on a DOX diet.

To address the potential mechanisms of tumor regression, we used histological analysis and flow cytometry to examine MIX 10:1 and wt tumors a week after the mice were placed on a DOX diet. Flow cytometry for T cells indicated a small number of infiltrated cells and no significant difference between MC38 and the MIX 10:1 tumors (**Fig. 2D**). A similar phenotype was observed for NK cells (Fig. S3). However, a massive amount of CD11b macrophages was detected in the MIX 10:1 tumors in some tumor specimens, reaching almost 40% of the total tumor cells (**Fig. 2E**). A similar number of infiltrated macrophages was observed in MIX 10:1 tumors isolated from nude mice (**Fig. 2F**). Immunohistochemical analyses of the macrophage marker CD68 indicated clusters of macrophages within the tumor tissue in both C57BL/6 and nude mice (**Fig. 2G**). Neutrophils promote an anti-cancer response by direct killing and by promoting adaptive T-cell responses (Granot, 2019). Using Ly6G as a marker, we assessed the presence of neutrophils in the TME. Similar to macrophages, neutrophils were enriched in MIX 10:1 tumor tissues in both C57BL/6 and nude mice (**Fig. 2H**, 2J). Immunohistochemistry revealed that neutrophils were homogenously dispersed in the tumor tissue (**Fig. 2I**). These data suggest robust infiltration of innate immune cells into the TME, primarily macrophages and neutrophils. This infiltration coincided with regression of tumors a few days after suppression of KDELR2 in as little as 10% of a small subset of tumor cells. CCL2 is a potent chemoattractant for multiple immune cells, including monocytes, macrophages (Balkwill, 2003; Zhang *et al*, 2010) and neutrophils (Johnston *et al*, 1999). Since CCL2 levels were enriched in the secretome of Tet-On shKDELR2, we checked whether this chemokine accumulated in the TME of MIX 10:1 tumors. Immunofluorescence analysis for CCL2 indicated an increase in the TME of MIX10:1 tumors (**Fig. 2K**), suggesting that this chemokine participates in inflammatory conditions that eventually lead to the regression of MIX 10:1 MC38 tumors when KDELR2 is targeted.

### Suppression of KDELR2 promotes phagocytosis of tumor cells, macrophage chemotaxis and polarization to M1 *in vitro*

At least three subtypes of tumor-associated macrophages exist (Dunsmore *et al*, 2024), of which M1 macrophages directly and indirectly promote anticancer immune responses (Mantovani *et al*, 2017; Sica *et al*, 2008). To assess whether the suppression of KDELR2 in tumor cells is sufficient to affect macrophage function, we generated bone marrow-derived macrophages (BMDM), polarized them to M1 with IFNγ+LPS, and analyzed their chemotaxis to supernatants of MC38 Tet-On shKDELR2 cells treated or not with DOX for 48 h. Using Boyden chambers, we found a strong chemotaxis of M1 BMDM in conditioned media collected from DOX-treated cells (**Fig. 3A**). We then tested whether the supernatants of shKDELR2-suppressed cells affect macrophage polarization. To this end, naïve BMDM were incubated for 24 h with conditioned media of Tet-On shKDELR2 pretreated with DOX. Based on the M1 and M2 markers iNOS and Arginase 1, supernatants collected from DOX-treated Tet-On shKDELR2 cells conferred an M1 phenotype comparable to the addition of IFNγ+LPS (**Fig. 3B**). No M2 macrophages were observed (**Fig. 3C**). We then examined whether the suppression of KDELR2 promotes the opsonization of tumor cells by M1 macrophages. We assayed this possibility by flow cytometry in which M1 polarized BMDM were labeled with Alexa647-anti-F4/80 (red) and mixed at a 1:5 ratio with CFSE-labeled Tet-On shKDELR2 MC38 cells (green) treated or not with DOX for 24 h. Only in the presence of DOX was a double-positive population detected, indicative of macrophage/tumor physical interactions (**Fig. 3D**). Microscopic analyses of the double-labelled population showed the presence of tumor cells either adhering to macrophages or actively undergoing phagocytosis (**Fig. 3E**). The secretome of Tet-On shKDELR2 MC38 cells was enriched in both KDEL and non-KDEL proteins. To determine whether the polarizing activity of the supernatants required KDEL proteins, we used an anti-KDEL antibody to deplete the supernatant of KDEL proteins. The anti-GFP antibody was used as an irrelevant negative control. Using three separate BMDM batches, the removal of KDEL proteins reduced polarization efficiency by more than 50% (**Fig. 3F**). We conclude that KDELR2 inhibition enables the polarization of macrophages into M1, promotes their chemoattraction, and opsonizes tumor cells for physical interaction with M1 macrophages. This was brought about by cooperation between KDEL and non-KDEL proteins.

**Figure 3:**
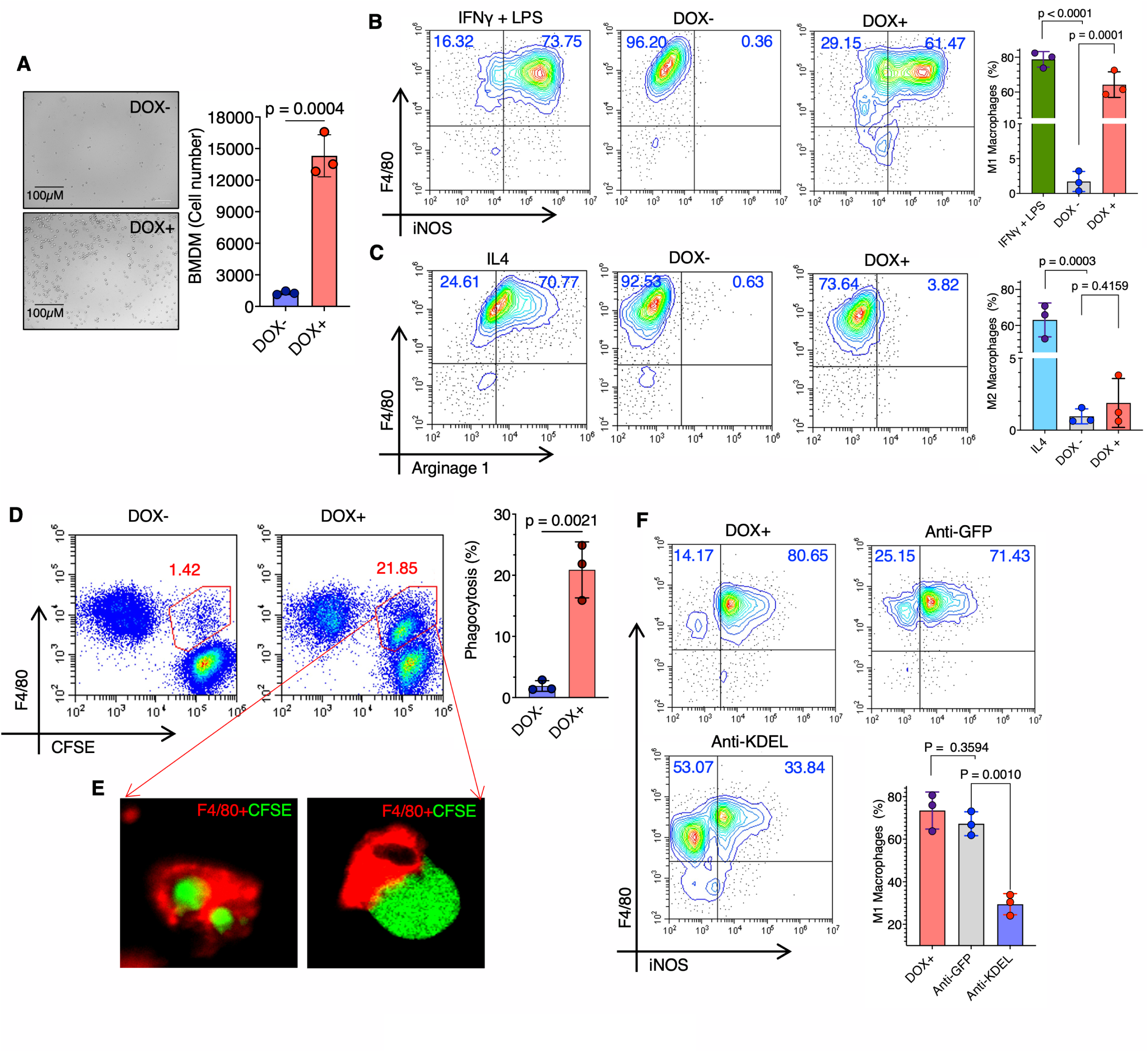
Suppression of KDELR2 promotes macrophage chemotaxis, polarization to M1 and opsonization of tumor cells. **A.** BMDM were generated by M-CSF, polarized to M1 by LPS+IFNγ and incubated with conditioned media of Tet-On shKDELR2 with and without DOX in a Boyden chamber. Chemotaxis was calculated by counting the cells attached to the separating filter. Shown is a representative image of the filter and quantification. **B** and **C.** Naïve BMDM were generated and incubated for 24 h with supernatants of MC38 or Tet-On KDELR2 cells that were pre-incubated with DOX. Cells were stained for F4/80, permeabilized and stained for intracellular iNOS (M1) or arginase 1 (M2). IFNγ or IL4 were used for positive controls of M1 and M2, respectively. Three independent repetitions were performed with similar outcomes. **D.** BMDM were polarized to M1, labeled with Alexa647-anti-F4/80 and mixed in a 1:5 ratio with CFSE labeled Tet-On shKDELR2 MC38 cells treated, or not with DOX for 24 h. Double positive cells were sorted, cyto-spin and imaged by epifluorescence. Adducts of macrophages and tumor cells were observed (**E**). **F.** Naïve BMDM were incubated with supernatants of Tet-On shKDELR2 preincubated with anti-KDEL or anti-GFP. Shown is a representative flow cytometry for M1 polarization of three.

### Suppression of KDELR2 by a lipid nanoparticle delivery of siRNA causes tumor regression

Because the suppression of KDELR2 in as little as 10% of tumor cells was sufficient to remodel the TME in a profound way, we reasoned that this approach could be recapitulated by conventional nano-delivery systems of siRNA, which are estimated to deliver their payloads at a similar efficiency (Fulton & Najahi-Missaoui, 2023; Silva-Cazares *et al*, 2020). To this end, we constructed siRNA-loaded LNPs 100 nm-300 nm in diameter with a zeta potential of 50 mV (Fig. S4). These positively charged particles were too large to extravasate easily from the injection site. Previous analyses have demonstrated knockdown efficiency for a few days *in vivo* (Nicolescu *et al*, 2022; Sun & Lu, 2023). To assess the uptake by MC38 cells, LNPs were labeled with Alexa647. Within 4 h of incubation with MC38 cells that express nanoluc-KDEL, most cells were labeled and the LNPs were observed intracellularly (**Fig. 4A**), most likely in endocytic compartments. qPCR analysis confirmed the reduction in KDELR2 mRNA levels (**Fig. 4B**) and an increase in luciferase activity in the supernatant (**Fig. 4C**), indicating the functional suppression of KDELR2 by the LNPs. We then examined the effects of the LNPs on tumor growth. C57BL/6 mice were inoculated with MC38, which stably expressed firefly luciferase (MC38-ffLuc). When tumors became palpable (approximately 500 mm^3^), 70 µL of LNPs with siRNA loaded with KDELR2 or a scrambled sequence was injected directly into the tumor. Shown for the individual mice, tumor growth was inhibited by the LNP carrying siKDELR2, but continued to grow in the scrambled siRNA control group (**Fig. 4D**, 4E). We then repeated the experiment, removing the tumors after 20 days. Flow cytometry analysis of tumor tissues indicated the enrichment of macrophages (**Fig. 4F**). Analysis of macrophage polarization indicated predominantly M1 macrophages (**Fig. 4G**), whereas macrophages in the scrambled control treatment were mostly M2 macrophages (**Fig. 4H**). Neutrophils were also observed at higher percentages in siKDELR2 treated tumors (**Fig. 4I**). Histological analysis of tumor sections showed the presence of macrophage clusters (**Fig. 4J**). CCL2 was abundant in tumors in which KDELR2 expression was suppressed (**Fig. 4K**). qPCR analyses of bulk RNA isolated from tumor tissues indicated an increase in CCL2 and TSP-1 mRNAs levels (**Fig. 4L**), suggesting enhanced *de novo* production 20 days after the initiation of KDELR2 inhibition.

**Figure 4:**
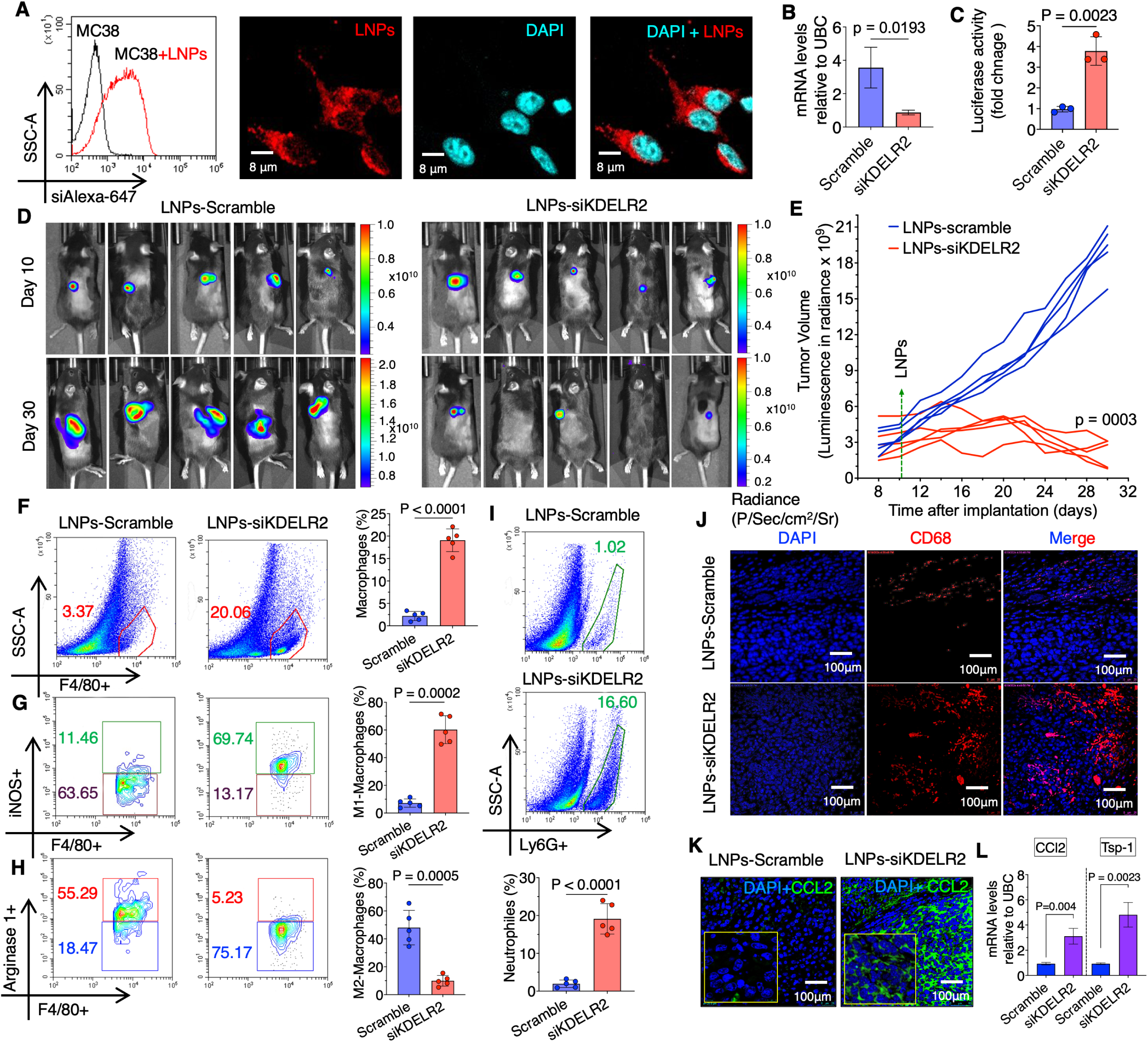
Suppression of KDELR2 by a lipid nanoparticle delivery of siRNA causes regression of MC38 tumors. **A.** MC38 cells were incubated with LNPs loaded with siRNA to KDELR2 and labeled with Alexa647. Shown by flow cytometry, most MC38 cells have taken up the nano particles. **B and C.** siKDELR2 LNPs suppress mRNA expression and promote luciferase activity in the supernatants of MC38/Nanoluc-KDEL cells. Shown is a represesentative experiment performed in three technical eplicates. C57BL/6 were inoculated with MC38-ffLuc cells. When tumors became palpable, LNPs carrying siRNA to KDELR2 or a scramble sequence control were injected into the tumor and growth was followed for an additional three weeks by total body bioluminiscence (N=5). Shown is the bioluminescence signal of every individual mouse (**D**) and averages (**E**). Tumors were extracted and analyzed by flow cytometry for total macrophages (**F**), M1 (**G**), M2 (**H**) neutrophils (**I**) and by immunohistochemistry for macrophages (**J**). **K.** Immunohistochemistry for CCL2. **L**. Total RNA was extracted from five tumors treated with siKDELR2 or a scrambled control. qPCR analysis shows an elevated level of CCL2 and TSP-1 mRNA.

### Suppression of KDELR2 causes the regression of B16F10 melanoma

MC38 is an antigenic tumor that is efficiently curtailed by T cells. B16F10 melanoma cells establish aggressive tumors that are poorly recognized by T cells, owing to low MHC class I expression (Lennicke *et al*, 2017) and high expression of PD-L1 (Basher *et al*, 2020). To examine whether KDELR2 inhibition by LNPs affects the growth of non-immunogenic tumors, C57BL/6 mice were challenged subcutaneously with B16F10 cells that stably expressed firefly luciferase. Fifteen days after implantation, when tumors were readily observed, 70 µL of LNPs carrying siRNA to KDELR2 or a scrambled sequence were injected into the tumor. Tumor growth was monitored using bioluminescence. Injection of KDELR2 siRNA LNPs resulted in retarded growth followed by regression, while the tumors continuously progressed in the control group. This was demonstrated in images of individual mice taken before LNP injection and at day 40 post-implantation, before euthanasia (**Fig. 5A**). The bioluminescence signal of each mouse over time is shown in **Fig. 5B**. A separate group of mice was used for TME analyses. Tumors were isolated at day 30 post-implantation, 15 days after the administration of LNPs, and analyzed by flow cytometry for macrophages and histology. Similar to what was observed for MC38, macrophage abundance increased in the TME upon KDELR2 suppression (**Fig. 5C**), and almost all exhibited an M1 phenotype (**Fig. 5D**). The predominant M2 phenotype was observed in the control group (**Fig. 5E**). Histological analysis showed clusters of macrophages in the siKDELR2 group (**Fig. 5F**). We conclude that remodeling of the TME in response to KDELR2 suppression is similar for hot and cold tumors.

**Figure 5:**
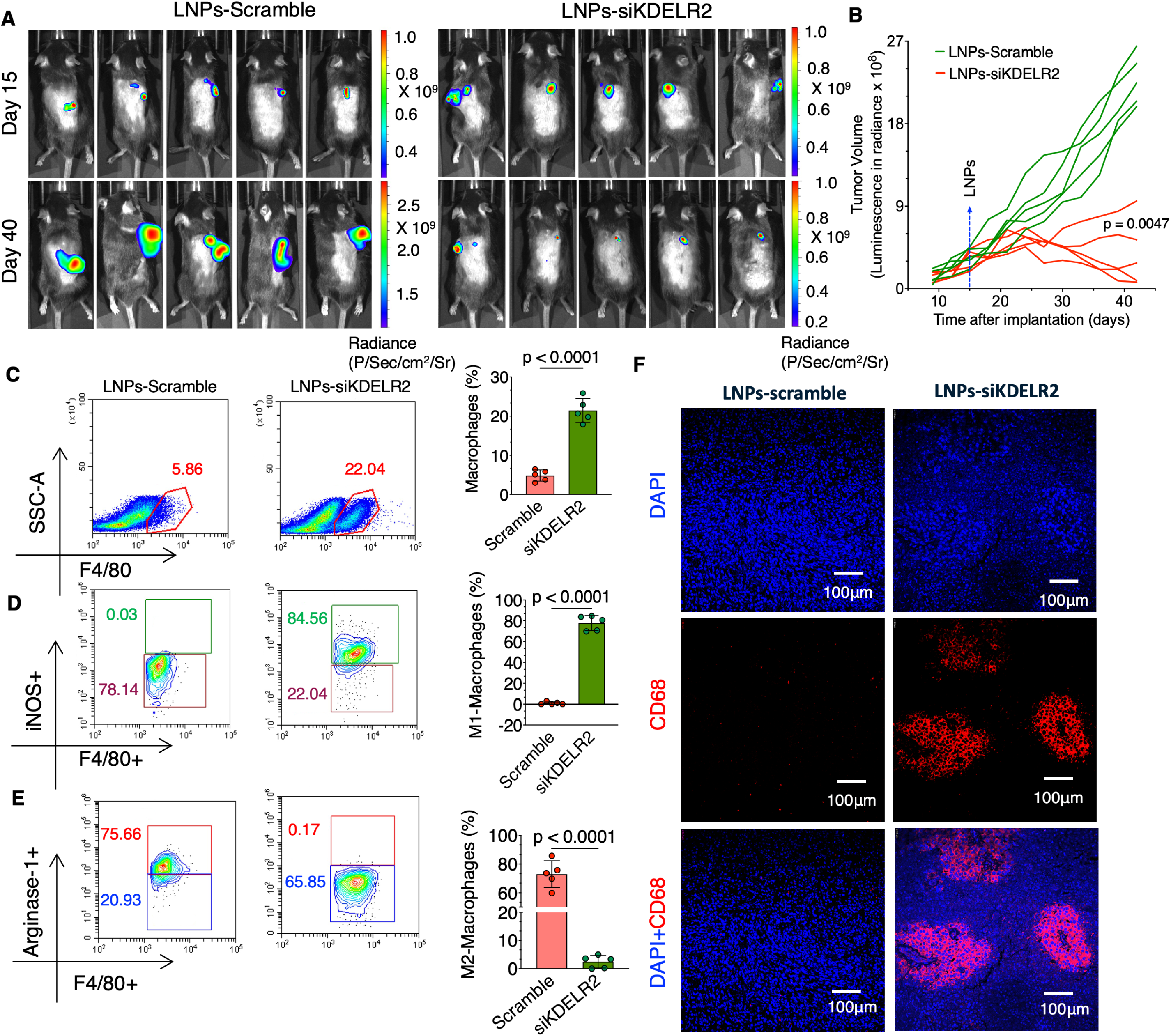
Suppression of KDELR2 by a lipid nanoparticle delivery of siRNA causes regression of B16F10 tumors. **A.** C57BL/6 males were inoculated with B16F10-ffLuc cells. When tumors became palpable, LNPs carrying siRNA to KDELR2 or a scramble sequence control were injected into the tumor and growth was followed by total body bioluminiscence (N=5). Shown is the bioluminescence signal of every individual mouse (**A**) and averages (**B**). Tumors were extracted and analyzed by flow cytometry for total macrophages (**C**), M1 (**D**) and M2 (**E**). **F.** Immunohistochemistry for macrophages of representative tumors from the two cohorts.

### Suppression of KDELR2 causes the regression of orthotopically implanted tumors

We next wondered whether tumors growing in tissues other than the skin would also exhibit infiltration of innate immune cells after KDELR2 suppression. To address this, we stably expressed firefly luciferase in an E0771 triple-negative breast cancer cell line. The cells were implanted into the mammary fat pads of C57BL/6 female mice. When reaching a size of 300-500 mm^3^, 70 µL of LNPs carrying siRNA to KDELR2 or a scrambled sequence were injected into the tumor. While the control tumors continued to grow, the E0771 tumors treated with siNA against KDELR2 grew slower and then started to regress (**Fig. 6A**). Of note, regression was slower than that for MC38 or B16F10 tumors. Moreover, 62 days after inoculation (47 days after LNP injection), the tumors were still detected (**Fig. 6B**). When analyzed histologically 30 days after administration of LNPs, infiltration of macrophages was observed, however, clusters were not observed (**Fig. 6C**). Flow cytometry estimated the percentage of macrophages to be approximately 15% (**Fig. 6D**). As observed for MC38 and B16F10 tumors, infiltrated macrophages were polarized primarily to M1 after KDELR2 silencing (**Fig. 6E**) and to M2 in the control group (**Fig. 6F**). In contrast to MC38 tumors, T cells were enriched in primary orthotopically implanted E0771 tumors (**Fig. 6G**), and most were CD8-positive (**Fig. 6F**). CCL2 was enriched in siKDELR2 tumors (**Fig. 6I**). We conclude that inhibition of KDELR2 remodels the TME to inhibit tumor growth in the mammary fat pad.

**Figure 6:**
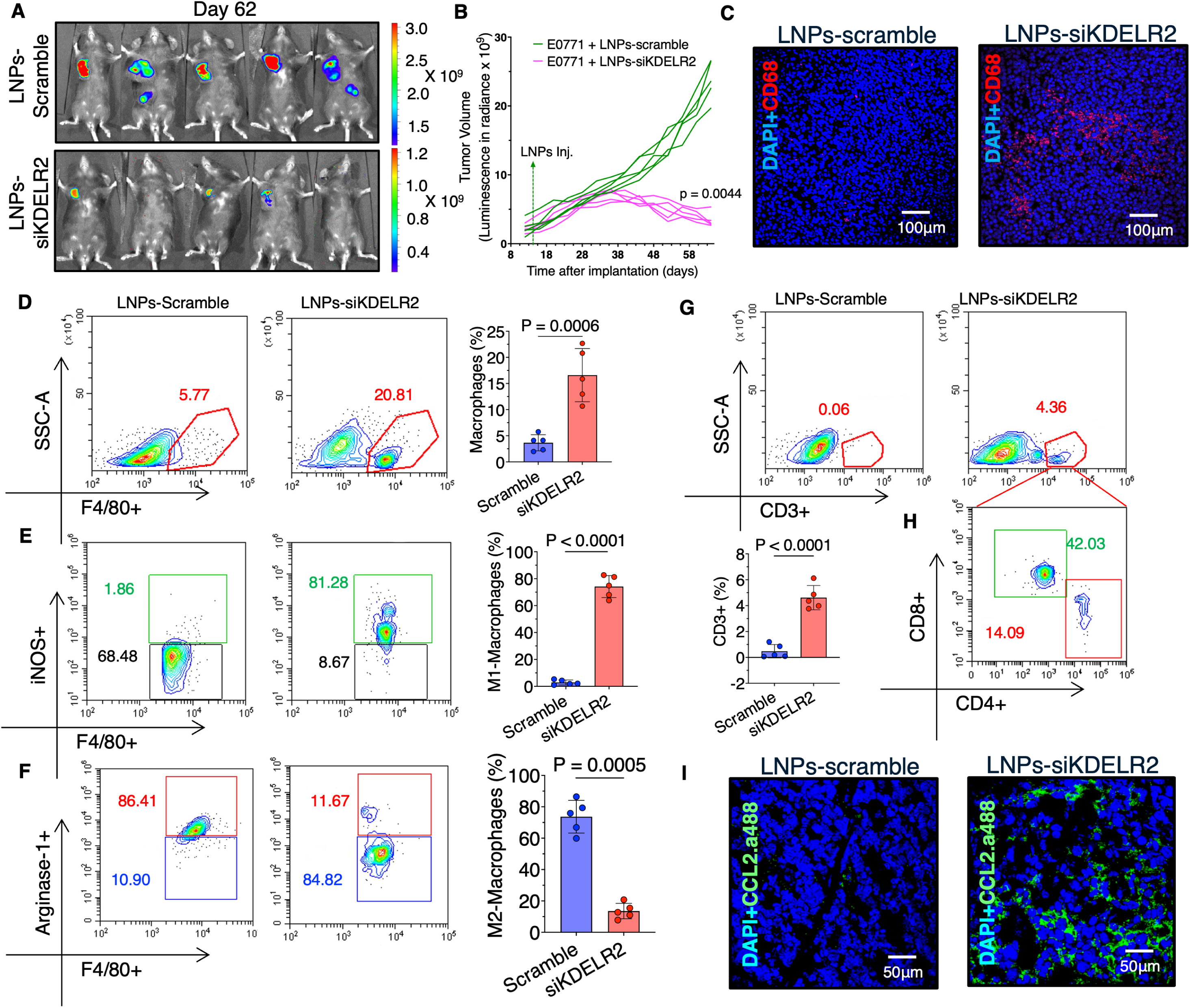
Suppression of KDELR2 by a lipid nanoparticle delivery of siRNA causes regression of orthotopic breast cancer tumors. **A.** C57BL/6 males were inoculated into the mammary fat pad with E0771-ffLuc cells. When tumors became palpable, LNPs carrying siRNA to KDELR2 or a scramble sequence control were injected into the tumor and growth was followed by total body bioluminiscence (N=5). Shown is the bioluminescence signal of every individual mouse (**A**) and averages (**B**). **C.** Tumors were extracted and analyzed by imunohostchemistry for macrophages (CD68). D. Flow cytometry analyses of the tumos shows an increase in macrophages (**D**) and polarization into M1 (**E**), while the macrophages were plorized into M2 in the scrambled control (**F**). **G and H**. Infiltrated T cells were mostly CD8-positive. **I**. imunohostchemistry for CCL2 demonstrates higher levels in the tumors treated with siKDELR2.

### Regression of KDELR2-suppressed tumors reduces tumor growth upon rechallenge, associated with enhanced T cell infiltration

T cells are essential for maintaining immunosurveillance and protection from tumor recurrence (van der Leun *et al*, 2020). To address whether ICD conditions induce T cell activation, we examined the growth of MC38 cells upon a second challenge after regression of KDELR2-silenced tumors. C57BL/6 mice were subcutaneously inoculated with MIX 10:1 or Tet-On shKDELR2 cells. Tumors were allowed to develop to 200 mm^3^, and then the mice were placed on a DOX diet until the tumors were no longer observed. Two weeks later, while still on the DOX diet, a second challenge with MC38 cells was performed. To ensure that the emerging tumors were not derived from dormant primary tumor cells, the second challenge was performed with MC38-ffLuc, and tumor growth was assessed by whole-body bioluminescence. Re-emerging tumors were confined to the injection location, without evidence of metastasis. When compared to the growth of MC38 cells in naive mice (blue lines), a significant decrease in tumorigenicity was observed (**Fig. 7A**, compare green to blue lines). Shown for individual mice, a stronger inhibitory effect was seen when only the Tet-On shKDELR2 were used for the first challenge (**Fig. 7A**, red lines), suggesting that inhibition of KDELR2 has a “dose” dependent effect against a rechallenge. Analysis of the tumors 34 days after rechallenge of C57BL/6 mice indicated the presence of T cells (**Fig. 7B**, 7C) composed of both CD4 and CD8 subtypes (**Fig. 7B**, 7D), consistent with previous studies showing a crucial role of T cells in controlling MC38 growth (Gong *et al*, 1997). Similar to what had been for the primary MC38 MIX 10:1 tumors, the rechallenged MC38 tumors were enriched for macrophages, predominantly M1 macrophages (Fig. S5), without any additional immune stimulation. To assess whether the retardation in growth was dependent on T cells, we repeated the experiment in nude mice. In this host, MC38-ffLuc growth was not significantly affected by regression of Tet-On shKDELR2 cells (**Fig. 7E**). These data suggest that the regression of tumors by the inhibition of KDELR2 initiates a systemic anti-cancer response that may protect against tumor dissemination. To address the possibility of T cell priming during the regression of primary tumors, we generated MC38 cells that express SIINFEKL, an ovalbumin-derived immunodominant H-2K^b^ MHC class I-restricted epitope. This epitope is recognized by T cells in OT-I TCR transgenic mice (Kurts *et al*, 1996). We purposely expressed the SIINFEKL peptide using the PresentER vector, which deposits the minimal epitope directly in the ER, obviating proteasomal degradation and TAP-dependent transport (Gejman *et al*, 2018) and avoiding confounding issues of stress-related effects on the MHC class I presentation machinery. MC38-SIINFEKL cells were mixed with Tet-On shKDELR2 cells at a 10:1 ratio and subcutaneously implanted into C57BL/6 mice. When the tumors were palpable, the mice were fed a DOX diet for a week. Before complete regression of the primary tumor, CFSE-labeled naïve CD8+ T cells isolated from the spleens of OT-I mice were adoptively transferred. Three days later, while still on the DOX diet, the mice were rechallenged on the other flank with either wt MC38 or MC38-SIINFEKL. One week later, the tumors were removed and analyzed by flow cytometry (**Fig. 7F**). While OT-I T cells were not evident in any of the analyzed wild-type MC38 tumors, we detected OT-I T cells at a frequency of approximately 3% of the total MC38-SIINFEKL tumor cells (**Fig. 7G**). Proliferation was observed with CFSE dilution, as shown for the two different tumors (**Fig. 7H**). This indicates a priming of tumor-recognizing T cells without active vaccination.

**Figure 7:**
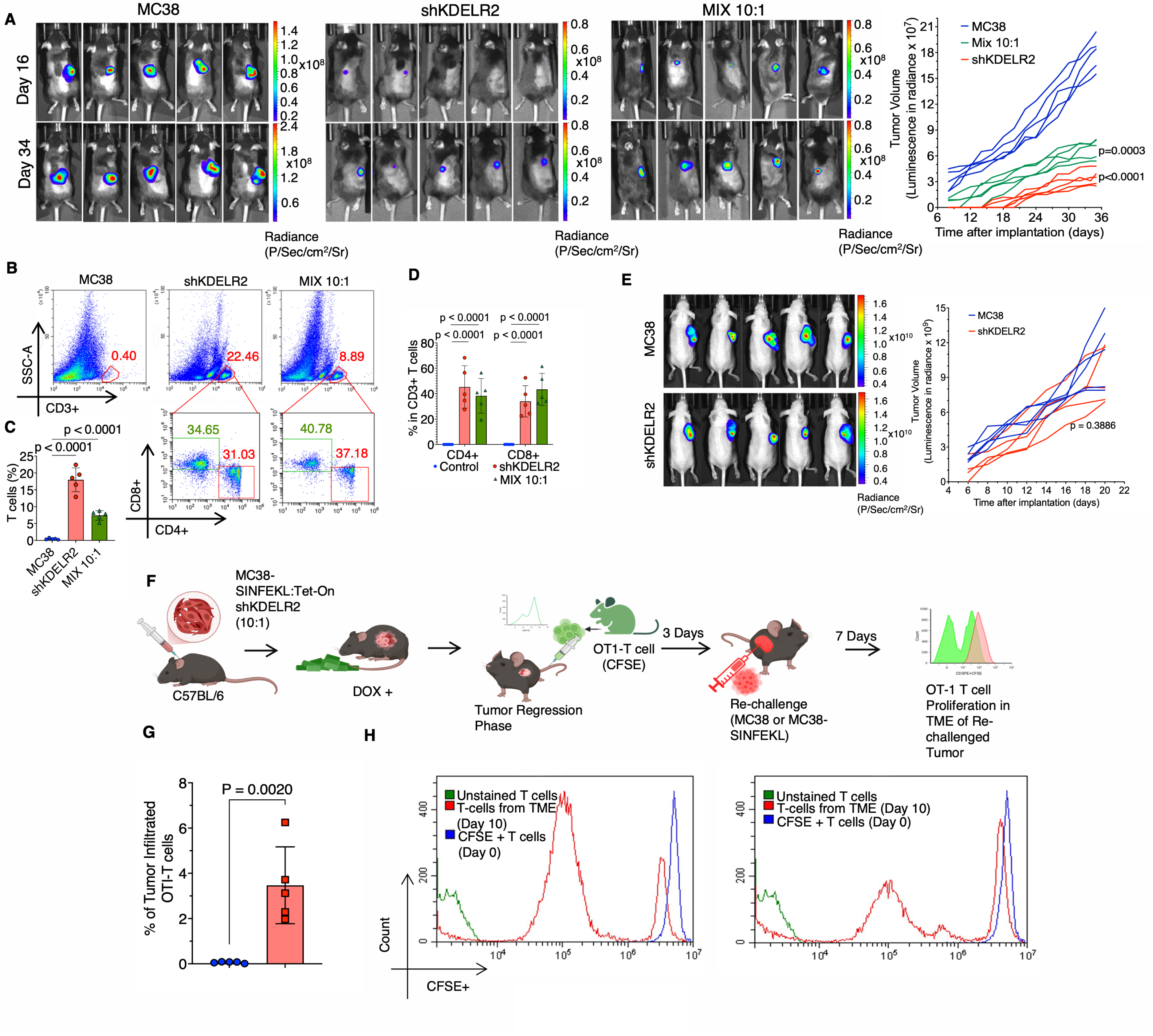
Suppression of KDELR2 induces T cell priming and a protective T cell response. **A.** C57BL/6 mice were implanted with MIX 10:1 or Tet-On KDELR2 cells. Tumors were allowed to progress to 800 mm^3^, then mice were placed on a DOX diet until tumors were no longer observed. Two weeks later, naive mice (blue lines) or mice in which tumors had been regressed were challenged with MC38-ffLuc cells and followed up to 34 days. Shown are the data for five mice in each group. **B**. MC38-ffLuc tumors from C57BL/6 were isolated and analyzed by flow cytometry for total T cells and CD4 and CD8 distribution. Total T cells are quantified in (**C**) and CD4/CD8 distribution in (**D**). **E.** Nude mice were treated as in A. Shown is the similar growth of MC38-ffLuc cells following tumor regression. **F.** C57BL/6 mice were challenged s.q. with a MIX 10:1 of MC38 that express SIINFEKL and MC38 Tet-On shKDELR2. Tumors were allowed to develop and then mice were put on a DOX diet. After a week naive OT-I T cells were adoptively transferred. Three days later mice were challenged with either wt MC38 or MC38 SIINFEKL. Seven days later, while mice are on a DOX diet, tumors were isolated and analyzed by flow cytometry for OT-I T cell proliferation. **G**. Shown are the levels of infiltrated OT-I cells in wt and SIINFEKL-expressing tumors, and (**H**) CFSE dilution pattern of OT-I T cells of two MC38 SIINFEKL tumors.

## Discussion

Except for platelets, in which PDI proteins are stored in secretory granules and secreted upon activation (Chen *et al*, 1995), in most cells, KDEL proteins are released in an uncontrolled manner, often due to ER stress (Wires *et al*, 2021). This indicates that, in most cells, the retention of KDEL proteins in the ER is inefficient and subject to leakage. Three potential mechanisms have been ascribed to the secretion of KDEL proteins: (i) secretion due to saturation of KDEL receptors (Trychta *et al*., 2018); (ii) inhibition of the interaction of the KDEL sequence with the cognate receptors (Li *et al*, 2015); and (iii) trafficking of the KDEL receptors to the plasma membrane, escorting their cargo for secretion (Wires *et al*., 2021). While these mechanisms probably operate in tandem, our data support a near saturation of the KDEL-KDELR interactions in MC38 cells (**Fig. 1**). For both KDELR1 and KDELR2, downregulation was sufficient to induce the release of KDEL proteins.

The relative abundance of the KDELR isoforms varies among cells (Bauer *et al*, 2020). We decided to silence each KDELR isoform separately to determine the dominant isoform in MC38 cells. Our data show that KDELR2 is the dominant gatekeeper isoform in this cell type, similar to what has been found in various human cells (Meng *et al*, 2022; Trychta *et al*., 2018). While not tested here, a standing question is whether the targeting of KDELR isoforms will provide stronger ICD conditions than the silencing of only one isoform. For optimal immunomodulatory bystander effects, a balance between cell viability and secretion should be considered. Most MC38 cells survived *in vitro* for up to four days after the initiation of KDELR2 knockdown (**Fig. 1**, **Fig. S1**). The silencing of the three KDELRs together might compromise cell viability faster, which would be insufficient to allow the accumulation of DAMPs in the TME. Defining the best combination and timing of KDELR silencing should be performed to optimize TME remodeling. The secretome of MC38 shKDELR2 cells was enriched in KDEL and non-KDEL proteins. This is not surprising, as KDEL proteins, primarily those engaged in protein folding, are required for the retention of non-KDEL proteins. One of the proteins that caught our attention was TSP-1, which was enriched in the KDELR2 secretome by more than four-fold. TSP-1 is retained in the ER in a PDIA3-dependent manner (Hecht *et al*, 2001; Hellewell *et al*, 2022). Once released, TSP-1 interacts with macrophages and promotes the secretion of IL-1β, a strong pro-inflammatory cytokine (Stein *et al*, 2016). It is therefore feasible that the modulation of the TME by KDELR2 inhibition is a result of the cooperative activity of the KDEL and non-KDEL proteins. The supernatants of KDELR2-inhibited MC38 cells conferred macrophage polarization (**Fig. 3**). This attribute has been ascribed to proinflammatory cytokines such as TNF-α, IL-1β, IL-6, and IL-12 (Sica & Mantovani, 2012). KDEL proteins have not been documented to directly affect polarization. Therefore, it was unexpected that the depletion of KDEL proteins from the supernatants would reduce the polarization efficiency (**Fig. 3**). This can be due to the action of KDEL proteins directly on macrophages, or, most likely, that KDEL proteins physically interact with cytokines and upon depletion, a portion of the cytokines is also removed. This signifies that KDEL proteins with chaperone activity, such as Bip, gp94 and calreticulin, when released *in situ*, may protect proinflammatory cytokines and promote their activity. We envision a positive feedback loop in which a small number of tumor cells release KDEL and non-KDEL proteins that initiate the infiltration of innate immune cells, macrophages, and neutrophils. Once in the TME, these immune cells are activated, polarized, and invigorate the infiltration by releasing chemokines and cytokines. This model may explain macrophage clusters. We assumed that neutrophils were not observed in the clusters owing to their higher mobility, allowing them to disperse through the tumor tissue. In support of this model, the TME is inundated with CCL2, a seminal chemokine for innate immune cells (Behfar *et al*, 2018).

The effect of KDELR2 inhibition in as little as 10% of the tumor cells was sufficient to cause complete regression of MC38 tumors. The growth of a second challenge was inhibited, but not prevented, even for antigenic tumors, such as MC38. The adoptive transfer of OT-I T cells indicates priming in a tumor antigen-dependent manner (**Fig. 7**). Priming of T cells requires an ordered lymphoid structure. Tertiary lymphoid tissues can be generated inside organs under prolonged chronic inflammatory conditions (Wang *et al*, 2023b). Although indirect evidence for T-cell priming within tumor tissues exists (Hildner *et al*, 2008; Sanchez-Paulete *et al*, 2017), it is unlikely that T-cell priming occurred *in situ*. A more likely scenario is the cross presentation of tumor antigens by presenting antigen cells at the local draining lymph nodes. This can be mediated by infiltrated M1 macrophages and/or dendritic cells, suggesting in-and-out trafficking. We propose that the enhanced permeability of the tumor tissue provoked by inflammatory conditions reprograms the TME to be non-permissive for tumor growth. These features seem to operate irrespective of T-cell antigenicity (**Figs. 4,5,6**). We propose that tumors that are unresectable but approachable for direct administration of RNA delivery systems, siRNA or antisense, which mildly respond to immunotherapy, would benefit most from this approach. This includes primary liver cancer or large liver metastases, which can be assessed using catheterization. Another possibility is metastatic large cell neuroendocrine carcinoma of the lung, which is a rare and aggressive type of lung cancer accessible by endoscopy (Andrini *et al*, 2022). We propose that the silencing of KDELRs can be leveraged to directly initiate CIC, overriding the need for therapy to induce ICD. To maximize the effect, a combination of KDELR inhibition with immune checkpoint inhibitors or CAR-T cells should be considered to sustain productive CIC.

## Materials and methods

### Sex as a biological variable

To avoid confounding immunological responses due to a gender mismatch, we decided to study the female cell lines MC38 and E0771 will be studied in female mice, and B16F10 melanoma cells in male mice.

### Cell lines

All cell lines were routinely checked for mycoplasma contamination by LookOut® Mycoplasma PCR Detection Kit (Millipore Sigma, cat# MP0035). MC38 cells were purchased from Kerafast (Shirley, MA). E0771 cells were purchased from ATCC. pcDNA3, which encodes the nanoluc-KDEL, was provided by Dr. Muhammad Mahameed (Hebrew University, Israel) and was used to stably express MC38. shRNA sequences targeting KDEL receptors were cloned into Tet-pLKO-puro (Addgene, cat# 21915). Oligonucleotide sequences were:

KDELR1: CCGGTGGTGTTCACTGCCCGATATCCTCGAGGATATCGGGCAGTGAACACCATTTTTG KDELR2: CCGGGACCATTCTCTACTGCGACTTCTCGAGAAGTCGCAGTAGAGAATGGTCTTTTTG KDELR3: CCGGGCAAGCTGTAAGCAGTCCAAACTCGAGTTTGGACTGCTTACAGCTTGCTTTTTG

The cells were infected with lentiviruses carrying shRNA sequences. Selection was performed using 1 µg/ml puromycin. Firefly luciferase was expressed in MC38 and E0771 cells by lentiviral infection, using pLX313-Firefly luciferase. B16F10 melanoma cells expressing firefly luciferase were kindly gifted by Dr. Serge Fuchs (U Penn, Philadelphia, PA, USA). All cells were maintained in DMEM supplemented with 10% fetal bovine serum (Corning, MT35010CV), 2 mM L-glutamine (Thermo Fisher Scientific, 25030081), 1% penicillin-streptomycin solution (Thermo Fisher Scientific, 15070063), and 1 mM sodium pyruvate (Thermo Fisher Scientific, 11360070).

### PCR and qPCR

Total RNA was extracted using QIAzol Lysis Reagent (Qiagen cat#79306), according to the manufacturer’s protocol. Total RNA (1 μg of total RNA was reverse-transcribed using the iScript cDNA Synthesis Kit (Bio-Rad, cat# 1708891) according to the manufacturer’s instructions. qPCR was performed using SsoAdvanced Universal SYBR Green Supermix (Bio-Rad, cat#1725270) on a CFX96 Touch Real-Time PCR machine (Bio-Rad). PCR for XBP1 mRNA splicing was performed using the REDExtract PCR ReadyMix (R4775, Sigma-Aldrich), as previously described (Drori *et al*, 2014). UBC was used as a normalizer.

Real Time Primers:

mKDELR1-F GATGGCAACCACGACACTTTCC, mKDELR1-R GGCAAGATAGCCACTGACTCCA mKDELR2-F CGATACCTTCCGAGTGGAGTTC, mKDELR2-R CCAGGTAGATGGAGAAGGTCCA mKDELR3-F GTGACTGGCCTTTCCTTTCT, mKDELR3-R CACCGGGAGGCTTAATTTCT mUBC-F CAGCCGTATATCTTCCCAGACT, mUBC-R CTCAGAGGGATGCCAGTAATCTA

### Secretome analyses by mass spectrometry

The cells were cultured in OptiMem supplemented with insulin/transferrin/selenium for 24 h in the presence or absence of 10 µM doxycycline. Cells were centrifuged, and the supernatants were concentrated with centricons with a cutoff of 3 kDa (Millipore # UFC500396), dried in SpeedVac, and reconstituted in 25 µL of 6 M urea/Tris buffer. The Protein samples were reduced with 120 mM DTT for 15 min at 55°C then alkylated with 500 mM iodoacetamide in the dark for 20 min at room temperature. The proteins were concentrated by overnight acetone precipitation at -20°C. Protein pellets were air dried for 10 min and dissolved in 40 μL 50mM tri-ethyl ammonium bicarbonate (TEAB) with 1 μg sequencing grade trypsin and incubated at 37 °C overnight. Digestion was terminated by adding TFA at a final concentration of 1%. The peptide samples were dried in a SpeedVac, reconstituted in 100 µL 0.1% formic acid, filtered through a 0.22 µm MilliporeSigma Ultrafree-MC Centrifugal Filter (#UFC30GVNB) before LC-MS/MS.

LC-MS/MS analysis: The samples were analyzed by LC-MS using a timsTof Pro2 instrument (Bruker) equipped with a NanoElute UHPLC system. A 5 µL aliquot of each digest was injected onto a ThermoScientific (0.5 x 5 mm) Acclaim Pepmap C_18_, 5-μm, trapping column. Liquid chromatographic elution was performed using a flow rate of 0.3 μl/min on a reverse phase column (ReproSil AQ C18, 75 μm x 150 mm, 1.9-μm 120-Å). Peptide elution was performed using a binary gradient of mobile phase A (0.1% formic acid) and mobile phase B (0.1% formic acid in acetonitrile). Each sample was analyzed using a linear gradient starting at 2% B at 0 min and increasing to 35% B in 30 min, followed by an increase to 90% B in 2 min, and then holding at 90% for 5 min before re-equilibration at 2% B. The electrospray voltage was 1500V. PASEF-DDA was used for peptide identification. MS1 scans were carried out with a resolution of 30,000, measuring masses between 100-1700 Da with 1/k0 values between 0.6 and 1.6 vs/cm^2^. MS2 scans between 100 and 1700 Da at a resolution of 30,000 were performed on the precursor between charge states of 2-5 with targeted intensities of 20000, an intensity threshold of 2500 au, and 10 PASEF MS/MS scans were performed in each cycle with cycle times of 1.2 sec. The peptides were fragmented using CID with an isolation window of 2 Da and collision energies ranging from 20 eV (1/k0 value of 0.6 Vs/cm2) to 59 eV (1/k0 value of 1.6 Vs/cm2). Dynamic exclusion was performed for 30 seconds.

Data analysis: LC-MS/MS data were searched against the mouse SwissProtKB database downloaded on 3-23-2022 (17,552 entries) using the program MSFragger v3.4. These searches were performed considering full tryptic peptides with no more than 2 missed cleavage sites. The MS1 and MS2 mass accuracies were set to 20 ppm, carbamidomethylation was considered a fixed modification, and oxidation of methionine and protein acetylation were considered variable modifications. The results were filtered using a reverse decoy database strategy with a percolator, and the PSM, Peptide, and protein FDR rates were set to 1%. Positively identified proteins were identified by a minimum of two peptides, with at least one of these being unique peptide identifications. Label-free quantification was performed using the MaxLFQ algorithm. The match between runs was enabled with a mass tolerance of 10 ppm, retention time shift of 5 min, and 1/k0 tolerance of 0.05. Quantitation was performed on unique peptides, only proteins identified by the two ions were quantified, and the LFQ intensities were normalized to the total peptide content for each LC-MS experiment. The LFQ intensities were used to determine the abundance ratios of the proteins across groups, and significance was determined using a T-T-derived p-value.

### Dual-Flank Xenograft-Tumor Model and Tissue Processing for Analysis

Two-month-old C57BL/6J female mice or female nude mice were challenged with 1×10^6^ MC38 cells suspended in 100 µL PBS in the left flank and modified Tet-On shKDELR2 MC38 cells in the right flank. Doxycycline was provided to the diet (Bio-Serv, cat# S3888). B16F10 ffLuc cells were inoculated in a manner similar to that for MC38 cells. Tumor volume was estimated from the length (L) and width (W), based on the formula: V = 1/6𝜋 x L x W x H. Following euthanasia, the tumors were carefully excised. A portion of each tumor was fixed in 4% paraformaldehyde for 24 h at room temperature and then transferred to ethanol for storage until further processing for histological analysis. The remaining tumor tissue was finely minced using sterile scissors and enzymatically digested in DMEM containing 0.2% collagenase (Sigma-Aldrich, Cat# SMB0070) at 37 °C with gentle agitation. After digestion, the cell suspension was filtered through a 70 µm cell strainer to remove debris and undigested fragments. The resulting single-cell suspension was used for downstream flow cytometry analysis.

### Immunoblotting

The cells were harvested by centrifugation (4 °C, 4000×g for 5 min) and washed with ice-cold PBS. Cell pellets were lysed in radioimmunoprecipitation assay (RIPA) buffer supplemented with protease inhibitors (Bimake, b14001) and phosphatase inhibitors (Bimake, b15001). Following vortexing for 10 min, the lysates were cleared by centrifugation (4 °C, 16,000×g for 15 min). The supernatant was separated and the protein content was quantified and mixed with reducing sample buffer, followed by boiling for 5 min. Samples were loaded on GenScript gradient SDS-PAGE gels, transferred to PVDF membranes (Immobilon-P) with eBlot (GenScript), detected by chemiluminescence using Immobilon^®^ Crescendo (Millipore, WBLUR0500), and imaged on a Bio-Rad ChemiDoc™ XR. Anti-nanoluc antibody (Promega, cat# N7000) was used according to the manufacturer’s instructions. p97 was used as the loading control (Novus Biologicals, cat# NB100-1558). HRP-conjugated goat anti-mouse and goat anti-rabbit secondary antibodies were purchased from Jackson ImmunoResearch Laboratories.

### Flow Cytometry Analysis

Tumor tissues were mechanically dissociated and passed through 70 µm cell strainers (Stellar Scientific, MD, USA) to obtain single-cell suspensions. For surface marker staining, the cells were incubated with fluorochrome-conjugated antibodies for 30 min at room temperature in the dark. The following antibodies were used: PerCP-conjugated anti-mouse CD45 (BioLegend, Cat# 103130), PerCP-conjugated anti-mouse/human CD11b (BioLegend, Cat# 101230), PE-conjugated anti-mouse CD3ε (STEMCELL Technologies, Cat# 130-116-289), APC-eFluor™ 780-conjugated anti-mouse CD4 (eBioscience/Thermo Fisher Scientific, Cat# 47-0042-82), and FITC-conjugated anti-mouse CD8a (eBioscience/Thermo Fisher Scientific, Cat# 11-0081-82). Appropriate isotype controls were obtained from BioLegend and BD Biosciences.

For intracellular staining, the cells were fixed and permeabilized using a BD Cytofix/Cytoperm™ Fixation/Permeabilization Kit (BD Biosciences, Cat# 554714), followed by staining with Alexa Fluor 488-conjugated anti-iNOS (Thermo Fisher Scientific, Cat# 53-5920-82) and APC-conjugated anti-Arginase-1 (Arg1) (Thermo Fisher Scientific, Cat# 17-3697-82).

Stained cells were acquired on a CytoFLEX S flow cytometer (Beckman Coulter) and analyzed using the CytExpert software. Gating strategies are provided in the supplemental supporting data (**Fig. S6, S7**).

### Immunohistochemistry and Immunofluorescence for Macrophage, Neutrophil & MCP1/CCL2 Analysis

Tumor tissues were fixed in 4% paraformaldehyde, dehydrated through an ethanol gradient, embedded in paraffin wax, and sectioned at 5 µm thickness. Paraffin-embedded sections were deparaffinized in xylene, rehydrated using descending ethanol concentrations, and rinsed in phosphate-buffered saline (PBS). Antigen retrieval was performed by boiling the sections in 10 mM sodium citrate buffer (pH 6.0) for 10 min.

After cooling, the sections were blocked for 1 h at room temperature in PBS containing 1% bovine serum albumin (BSA) and 2% goat serum. Slides were then incubated overnight at 4 °C with primary antibodies: anti-CD8 (add vendor and catalog number) and anti-MCP1/CCL2 (Thermo Fisher Scientific, Cat# MA5-17040).

For immunofluorescence, the slides were washed three times in PBS with 0.1% Tween-20 (PBST), followed by 1 h incubation at room temperature with fluorophore-conjugated secondary antibodies: Alexa Fluor® 488 (Jackson ImmunoResearch, Cat# 115-545-003) and Alexa Fluor™ 546 (Thermo Fisher Scientific, Cat# A-11081). After the final washes, sections were mounted using VECTASHIELD® PLUS Antifade Mounting Medium with DAPI (Vector Laboratories, Cat# H-2000). Imaging was performed using a Leica SP8 confocal microscope (Leica Microsystems), and images were analyzed using the LAS X software (Leica Application Suite X).

For chromogenic immunohistochemistry (IHC), adjacent sections were similarly processed for antigen retrieval and primary antibody incubation. After washing, the sections were incubated with HRP-conjugated secondary antibodies and developed using 3,3’-diaminobenzidine (DAB) substrate. Nuclei were counterstained with hematoxylin. The slides were visualized using an Olympus microscope equipped with differential interference contrast (DIC) optics (Olympus).

### Generation of BMDMs, Polarization Assays, and Chemotactic Response

BMDM were generated from femur bone marrow cells of 2-month-old female C57BL/6J mice. In brief, after flushing the bone marrow cells using a syringe and 23G needle, red blood cells were removed with ACK lysis buffer (cat# A1049201, Thermo Fisher, USA). After washing, the cells were cultured in a bone marrow culture medium (DMEM supplemented with 10% FBS, 1% penicillin-streptomycin, and 1 ng/mL M-CSF1) for six days. On day 7, BMDMs were washed twice with DMEM and treated with 30% v/v concentrated supernatants collected from shKDELR2-MC38 cells diluted in DMEM for 48 h. For positive controls, M1 polarization was induced using IFNγ (20 ng/mL), PeproTech®, Thermo Fisher, cat#AF-315-05-20UG, and LPS (240 ng/mL, Sigma-Aldrich, cat#SMB00704), while M2 polarization was induced using IL-4 (20 ng/mL, PeproTech®, Thermo Fisher, cat#214-14-20UG). The cells were analyzed by flow cytometry. M1 macrophages were identified as double-positive for anti-F4/80 and anti-iNOS staining, whereas M2 macrophages were identified as double-positive for anti-F4/80 and anti-arginase 1 staining. In a separate experiment, chemotaxis was evaluated using the Cell Migration/Chemotaxis Assay Kit from Abcam (cat# ab235673) following the manufacturer’s instructions. Briefly, BMDMs were seeded in the upper chamber of the assay insert, which contained a porous membrane. The lower chamber was filled with supernatant from shKDELR2-MC38 cells. The cells were incubated for 24 h. A positive control for the chemotaxis assay was included, as described in the kit protocol.

### Immunodepletion of KDEL-Tagged Proteins from Tumor Cell Supernatants and Functional Analysis in BMDMsKDEL-Depletion assay

The culture supernatants from both MC38 and shKDELR2-MC38 cells were incubated overnight at 4 °C with either anti-KDEL (Invitrogen, Thermo Fisher, cat# MA5-27581) or anti-GFP antibodies (Invitrogen, Thermo Fisher, cat# MA5-15256) bound to Protein A/G agarose beads (2 µg antibody per 20–30 µL beads for every 1 mL of supernatant). After incubation, the samples were centrifuged at 500 × g for 5 min at 4 °C to pellet the beads. The resulting supernatants were collected, passed through a 0.22 µm filter, and added to day 6 BMDM cultures for 48 h incubation. Following treatment, the cells were harvested and analyzed using flow cytometry. M1 macrophages were identified based on dual staining for F4/80 and iNOS, and their frequencies were compared to those of positive controls that had been treated with non-depleted (anti-GFP) supernatants.

### Phagocytosis Assay Using IFNγ/LPS-Activated BMDMs Co-cultured with CFSE-Labeled MC38 or shKDELR2-MC38 Cells

As outlined above, BMDMs were cultured in a bone marrow culture medium for six days. On day 7, IFN-γ (20 ng/mL) and LPS (240 ng/mL) were added to induce activation. After 48 h of stimulation, the cells were harvested and transferred to serum-starved medium for 12 h for phagocytosis assay.

Separately, 2 × 10⁶ MC38 or shKDELR2-MC38 knockout cells were harvested and labeled with CFSE (5 μM) for 10 min. Following BMDM harvesting, 2 × 10⁴ macrophages were co-cultured with equal numbers of CFSE-labeled MC38 or shKDELR2-MC38 cells for 2 h. After co-culture, cells were collected in FACS tubes, stained with anti-mouse F4/80 antibody, and analyzed for phagocytosis efficiency. Cells that were double positive for CFSE and F4/80 were sorted using a BD Aria III cell sorter and further examined using a Leica SP8 confocal microscope.

### Immune Memory Assessment in MC38-ffLuc Re-challenge Model

C57BL/6 mice were subcutaneously implanted with a mixture of Tet-on-shKDELR2 MC38 cells and parental MC38 cells at a 10:1 ratio and a separate group with Tet-on-shKDELR2 MC38. Upon tumor establishment, animals were maintained on a doxycycline-supplemented diet for two weeks to induce shRNA expression. The tumor volumes were monitored weekly using calipers. Once complete tumor regression was observed (volume approaching zero), doxycycline administration was discontinued, and the mice underwent a washout period of one week.

Following the washout, the mice were re-challenged with MC38 cells stably expressing firefly luciferase (MC38-ffLuc) via subcutaneous injection. A separate cohort of naïve C57BL/6 mice that received the same Luc-labeled MC38-ffLuc cells served as the control group. Tumor progression in both the experimental and control groups was assessed longitudinally using bioluminescence imaging (BLI) performed on the IVIS Spectrum system (PerkinElmer, USA).

Prior to imaging, the mice were injected intraperitoneally with 100 mg/kg D-luciferin. Anesthesia was induced and maintained using vaporized isoflurane (Abbott Laboratories, IL, USA) and the mice were positioned in the imaging chamber. Bioluminescent signals were captured 8 min post-injection, with a fixed exposure time of 60 s per mouse, throughout the duration of the study.

Quantitative analysis of the bioluminescence data was conducted using the Living Image software platform (PerkinElmer). Photon flux was quantified as the average radiance (photons/s/cm ²/sr) within a defined rectangular region of interest (ROI) encompassing the tumor-bearing area. Raw data were further processed and visualized using Newton module integration within the Amira platform and MATLAB (MathWorks, USA) to ensure consistency in image registration and signal normalization across the time points.

### Adoptive transfer of OT-I T cells

Naïve T cells were isolated from the spleen of OT-I mice using the EasySep™ Mouse T Cell Isolation Kit (Catalog # 19851). The cells were labeled with CFSE (Thermo Fisher, cat# C34554) according to the manufacturer’s instructions. A total of 10^7^ cells were suspended in 200 µL PBS and injected intravenously into tumor-bearing C57BL/6 mice.

### Formulation of siRNA LNPs and their Biophysical Characterization by Gel Retardation and DLS

ECO (MW ¼ 1023) at a stock concentration of 50 mM in ethanol was vortexed with siScramble duplex: sense 5′-GUUUCACACGUCGUACUCACU-3′ and antisense 5′- AGUGAGUACGACGUGUGAAAC-3,’ siKDELR2 sense: 5′-GACCAUUCUCUACUGCGACUU-3′; antisense: 5′-AAGUCGCAGUAGAGAAUGGUC-3’ at a stock concentration of 250 μM in nuclease-free water for 30 min to form nanoparticles at N/P = 8 with a final siRNA concentration of 100 nM. Helper lipids, beta-sitosterol (BCHOL) (Millipore Sigma), were added by combining an equivalent volume of a helper lipid with ECO at a concentration of 0.5 mM. Agarose gels (1% w/v) were prepared in Tris/borate/ethylenediaminetetraacetic acid (EDTA) buffer with ethidium bromide added for the gel retardation assay. A nanoparticle formulation (10 μL) was added to 2 μL of DNA gel-loading dye (6X, Life Technologies) and run at 100 V for 20 min. The gel was imaged using a ChemiDoc XRS system (Bio-Rad). The nanoparticle size and zeta potential were measured using dynamic light scattering on an Anton Paar Litesizer.

### LNP Treatment in MC38-ffLuc and B16F10-ffLuc Xenograft Models

When tumors derived from MC38-ffLuc or B16F10-ffLuc cells reached an average volume of ∼160 mm³, a single intratumoral injection of lipid nanoparticles (LNPs) containing either scrambled siRNA (control) or siRNA targeting KDELR2 (siKDELR2) was administered in a fixed volume of 60 µL. Tumor progression was subsequently monitored using bioluminescence imaging (BLI) to evaluate changes in tumor burden over time.

Imaging was continued until a marked reduction in luminescent signal was observed in the siKDELR2-treated group relative to the scramble control group, and until tumors in the control group reached volumes near the ethical endpoint (necrosis or ulceration).

In parallel, two additional cohorts for each tumor model underwent the same LNP treatment regimen. However, in these groups, tumors were harvested two weeks post-injection for immunological characterization, including analyses of tumor-infiltrating immune cells and cytokine profiling within the tumor microenvironment.

### Orthotopic Implantation and LNP Treatment in E0771-ffLuc Mammary Tumor Model

E0771-ffLuc cells were cultured under standard conditions and resuspended at a concentration of 5 × 10⁶ cells/mL in sterile PBS. To preserve the cell viability, the cell suspension was kept on ice and used for implantation within one h of preparation. For orthotopic tumor establishment, 100 µL of the cell suspension was injected into the mammary fat pad between the second and third mammary glands of female C57BL/6 mice using a 28.5-gauge needle.

Following tumor engraftment, bioluminescence imaging (BLI) was performed to monitor tumor development. Once the tumors reached a palpable volume of approximately 180 mm³, a single intratumoral injection of lipid nanoparticles (LNPs) containing either scrambled siRNA (control) or siKDELR2 was administered at a volume of 60 µL per tumor.

Tumor progression was monitored longitudinally via BLI to assess changes in the luminescence signal intensity, reflecting the treatment response. At the end of the study, tumors were harvested for comprehensive immunological analyses to evaluate the tumor microenvironment (TME), including immune cell infiltration and cytokine expression profiles.

### Statistical analysis

Statistical analyses were performed using R or GraphPad Prism software, version 10.0. The values are reported as the mean ± SD. Data were compared between groups using one-way analysis of variance (ANOVA) followed by Tukey’s multiple comparisons post-hoc test. The percentage of immune cell infiltration was visualized using a violin plot showing the median value, and comparisons between the two groups were performed using an unpaired t-test. P was set at p < 0.05. *In vivo* experiments were performed in a semi-blind fashion.

### Study Approval

Care of the experimental animals was performed in accordance with the institutional guidelines. The procedures were performed in accordance with the protocol approved by the IACUC of CWRU, protocol # 2023-0013.

### Data availability

All data including the secretome analyses are provided in the supplemental data.

### Author Contributions

Conceptualization: S.P.P., W.C.M., B.T.

Methodology: S.P.P., H.W., Z.R.L., B.T.

Formal analysis: S.P.P.

Investigation: S.P.P., B.T.

Resources: B.T., Z.R.L, B.W.

Data Curation: S.P.P., B.W.

Writing- original draft: S.P.P., W.C.M, B.T.

Supervision: B.T.

Funding acquisition: B.T., S.P.P.

## Acknowledgments

This research was supported by the National Cancer Institute 1R01CA299332-01 (BT) and The Elsa U. Pardee Foundation (SPP, BT). The timsT of Pro2 instrument was purchased from an NIH shared instrument grant (S10OD030398). The authors acknowledge the assistance of the Case Western Reserve University School of Medicine Light Microscopy Imaging Facility and support from the NIH grant S10OD02499601.

